# Shared and distinct transcriptomic cell types across neocortical areas

**DOI:** 10.1101/229542

**Authors:** Bosiljka Tasic, Zizhen Yao, Kimberly A. Smith, Lucas Graybuck, Thuc Nghi Nguyen, Darren Bertagnolli, Jeff Goldy, Emma Garren, Michael N. Economo, Sarada Viswanathan, Osnat Penn, Trygve Bakken, Vilas Menon, Jeremy Miller, Olivia Fong, Karla E. Hirokawa, Kanan Lathia, Christine Rimorin, Michael Tieu, Rachael Larsen, Tamara Casper, Eliza Barkan, Matthew Kroll, Seana Parry, Nadiya V. Shapovalova, Daniel Hirchstein, Julie Pendergraft, Tae Kyung Kim, Aaron Szafer, Nick Dee, Peter Groblewski, Ian Wickersham, Ali Cetin, Julie A. Harris, Boaz P. Levi, Susan M Sunkin, Linda Madisen, Tanya L. Daigle, Loren Looger, Amy Bernard, John Phillips, Ed Lein, Michael Hawrylycz, Karel Svoboda, Allan R. Jones, Christof Koch, Hongkui Zeng

## Abstract

Neocortex contains a multitude of cell types segregated into layers and functionally distinct regions. To investigate the diversity of cell types across the mouse neocortex, we analyzed 12,714 cells from the primary visual cortex (VISp), and 9,035 cells from the anterior lateral motor cortex (ALM) by deep single-cell RNA-sequencing (scRNA-seq), identifying 116 transcriptomic cell types. These two regions represent distant poles of the neocortex and perform distinct functions. We define 50 inhibitory transcriptomic cell types, all of which are shared across both cortical regions. In contrast, 49 of 52 excitatory transcriptomic types were found in either VISp or ALM, with only three present in both. By combining single cell RNA-seq and retrograde labeling, we demonstrate correspondence between excitatory transcriptomic types and their region-specific long-range target specificity. This study establishes a combined transcriptomic and projectional taxonomy of cortical cell types from functionally distinct regions of the mouse cortex.

## INTRODUCTION

The neocortex coordinates most flexible and learned behaviors. Based on cytoarchitectonic, neurochemical, connectional and functional studies, up to 180 distinct cortical areas have been identified in humans (Glasser et al., 2016) and dozens in rodents (Kolb, 1990; Ng et al., 2009). Cortical areas are often categorized as sensory, motor or associational, based on their connections with other brain areas. Each cortical area has a laminar structure (layers (L) 1-6). Most sensory areas contain L4, and they are termed 'granular', whereas motor and associational areas usually do not have L4, and are hence 'agranular'. In vertebrate evolution, cortex underwent disproportionally greater expansion in terms of the number of cells, layers and functional areas compared to the rest of the brain, coinciding with the acquisition of increasingly sophisticated cognitive functions (Jerison, 2009; Kaas, 2009; Man et al., 2013; Van Essen et al., 2016). Therefore, a systematic characterization of the structure and function of neocortex is essential for understanding neural mechanisms underlying cognition and behavior in animals and human.

Much effort has been devoted to understanding the extent of cortical cellular diversity and how the dynamics of cortical neural activity emerges from its structure – often referred to as the cortical circuit. Although cortical areas share some features including layered organization and certain connectivity patterns, neurons in different cortical areas show qualitatively different activity patterns that support different cortical computations. At one extreme, granular primary visual (VISp) and other sensory cortical areas process sensory information with millisecond time-scale dynamic (Cardin et al., 2010; Durand et al., 2016; Liu et al., 2009). Agranular frontal areas, including prefrontal cortex and motor cortex, such as the anterior lateral motor cortex (ALM) in mice, show slow dynamics related to short-term memory, deliberation, decision-making and planning (Chen et al., 2017; Guo et al., 2017; Guo et al., 2014). Categorizing cortical neurons into distinct types, and studying the roles of different types in the function of the circuit, is an essential step towards understanding how these different cortical circuits produce distinct computations (Svoboda and Li, 2017; Zeng and Sanes, 2017).

Previous studies, mainly in the rodent primary somatosensory, visual, and motor cortices, have identified multiple types of glutamatergic and GABAergic neurons (Molyneaux et al., 2007; Rudy et al., 2011). Recent large-scale neurophysiological and anatomical studies have attempted more comprehensive descriptions of cortical neuronal types, identifying dozens to hundreds of putative cell types (Jiang et al., 2015; Markram et al., 2015). An alternative approach to taxonomy relies on genome-wide gene expression by single-cell RNA-sequencing (scRNA-seq), in combination with unsupervised feature selection and cluster analysis (Li et al., 2017; Macosko et al., 2015; Pollen et al., 2015; Shekhar et al., 2016; Tasic et al., 2016; Zeisel et al., 2015). In a scRNA-seq study on mouse neocortical and hippocampal cells, 3,005 cells were profiled, leading to the identification of 29 neuronal transcriptomic types (Zeisel et al., 2015). We previously obtained scRNA-seq data from 1,679 cells from the mouse primary visual cortex (VISp) and classified them into 19 excitatory, 23 inhibitory, and 7 non-neuronal types (Tasic et al., 2016). Reconciling the morphological, neurophysiological and scRNA-seq data into a consensus picture of cortical neuronal types remains a major challenge.

A major advantage of scRNA-seq compared to other approaches to neuronal characterization is scalability. We leveraged it to conduct a large-scale scRNA-seq study of neocortical cell types with two goals in mind. First, to profile a sufficient number of VISp neurons to saturate the discovery of new transcriptomic cell types with our scRNA-seq approach. Second, to profile a different cortical area (ALM) at similar depth to test how cortical neuronal types vary across cortical areas with distinct functions.

Using SMART-Seq v4-based deep sequencing approach, we profiled more than 22,000 VISp and ALM cells isolated from a diverse set of Cre lines, as well as from retrograde labeling. Dimensionality reduction and clustering of the dataset gave rise to 116 transcriptomic types, 67 of which are shared between the two areas. Overall, this represents the most comprehensive cortical cell type classification to date. We find that interneuron and non-neuronal types are shared between the two cortical areas, whereas most excitatory neuron types are unique to each area. By combining retrograde labeling with scRNA-seq, we reveal clear correlation between excitatory neuron projection patterns and their transcriptomic identities. Finally, we provide correspondence to known cell types, and identify numerous genetic markers that can be used to build tools, including several transgenic recombinase lines presented here, to study the identified transcriptomic types in the future. Together with the accompanying paper (Economo et al., 2017), our study suggests a general principle: distinct computations in different cortical areas are based on their molecularly distinct projection neurons and their multi-regional connections.

## RESULTS

### Overall cell type taxonomy

Building on our previous pilot study (Tasic et al., 2016), we established a scRNA-seq pipeline with optimized, SMART-Seq v4 based methodology, standard operating procedures and stringent quality control (QC) criteria (**Extended Data Figs. 1 and 2**). Individual cells were isolated by fluorescence activated cell sorting (FACS) or manual sorting, cDNA was generated and amplified by SMART-Seq v4, and Nextera XT-tagmented libraries were sequenced on the Illumina HiSeq2500 platform (**Methods**). The major methodological differences between the previous pilot study and the current pipeline include the use of a N-methyl-D-glucamine (NMDG)-based artificial cerebrospinal fluid (ACSF) to improve neuronal survival (Ting et al., 2014) and replacement of Clontech’s SMARTer v1 with SMART-Seq v4 kit (**Methods**). SMART-Seq v4 is based on Smart-seq2 (Picelli et al., 2013), and increases gene-detection efficiency compared to SMARTer v1 (Ramskold et al., 2012) used in our previous study (Tasic et al., 2016). This allowed us to reduce the median sequencing depth from ~8.7 to ~2.6 million reads per cell while still detecting ~8,200 genes per cell (median) compared to ~7,800 previously (**Extended Data Fig. 2b**). This dataset represents the largest survey of single-cell transcriptomics data (57.18×10^9^ reads) from the neocortex in any species to date.

For clustering, we used 21,749 QC-qualified single-cell transcriptomes from VISp and ALM of adult mice (postnatal day P56±3) of both sexes in the congenic C57BL/6J background (**Extended Data Fig. 1a, Methods**). To cover neuronal cell types as comprehensively as possible while retaining layer-of-origin information, we employed layer-enriching dissections from ALM and VISp of *pan*-neuronal, *pan*-excitatory or *pan*-inhibitory recombinase lines crossed to recombinase reporters (referred to as the PAN collection) consisting of 11,153 cells (**Extended Data Fig. 1b, Extended Data Table 1**). To sample non-neuronal cells, we collected 816 reporter-negative cells (408 from ALM and VISp, each) from a *pan*-neuronal *Snap25-IRES-Cre* line. To compensate for cell type-specific survival biases and to collect rare types, we supplemented the PAN collection with 7,919 cells isolated from a variety of other recombinase lines, with or without layer-enriching dissections. These recombinase lines were selected to capture additional cellular diversity suggested by ongoing analysis of the PAN-collection data (**Extended Data Fig. 1b,e,f**). Finally, to investigate the correspondence between transcriptomic types and projection properties, 2,677 retrogradely labeled cells were sequenced.

To define transcriptomic types, single-cell transcriptomes were subjected to feature extraction and clustering (**Extended Data Fig. 2c, Methods**). To extract relevant transcriptomic features, we employed dimensionality reduction by weighted gene co-expression network analysis (WGCNA) (Langfelder and Horvath, 2008). We combined this approach with hierarchical and graph clustering in an iterative and bootstrapped manner (**Extended Data Fig. 2c). T**he output of this procedure is a co-clustering matrix: a cell-cell matrix that indicates the frequency with which any cell clusters with any other cell in 100 bootstrapped iterative clustering rounds (**Extended Data Fig. 3a**). Final transcriptomic cell types are defined by 'cutting' the co-clustering matrix to derive membership of each cell to a cluster (116 clusters total). Each cell’s membership also is tested post-clustering by classification algorithms to assign core vs. intermediate identity to cells (**Methods, Extended Data Fig. 2c)** (Tasic et al., 2016). Cells that are reliably classified to only one cluster (in >90 of 100 trials) are labeled “core” cells (19,195 cells), whereas cells that fall into more than one cluster are labeled “intermediate” cells. The vast majority of intermediate cells are assigned to only two clusters (2,542 of 2,554; 99.5%). The sequencing depth and gene detection per cluster are shown in **Extended Data Fig. 4.**

The identity of each cluster was assigned based on previously reported and newly discovered differentially expressed markers (**Extended Data Figs. 5 and 6)** and resulted in 52 glutamatergic, 50 GABAergic and 14 non-neuronal types (**Fig. 1**). These types correspond well to the 49 types from our previous study, with some boundaries between types shifted and better resolution provided in the current dataset (**Extended Data Fig. 7**). In addition, we now detect some rare types that were not sampled in the previous dataset. To determine if our sampling reached a plateau for cell type identification, we subsampled cells from the dataset and evaluated the stability of cluster identity with fewer cells. We find that for the vast majority of clusters, we sampled many more cells than needed to define them, whereas for a few (e.g., certain non-neuronal types, L5 PT and L6b types), additional sampling may reveal additional diversity (**Extended Data Fig. 8**).

**Figure 1.**
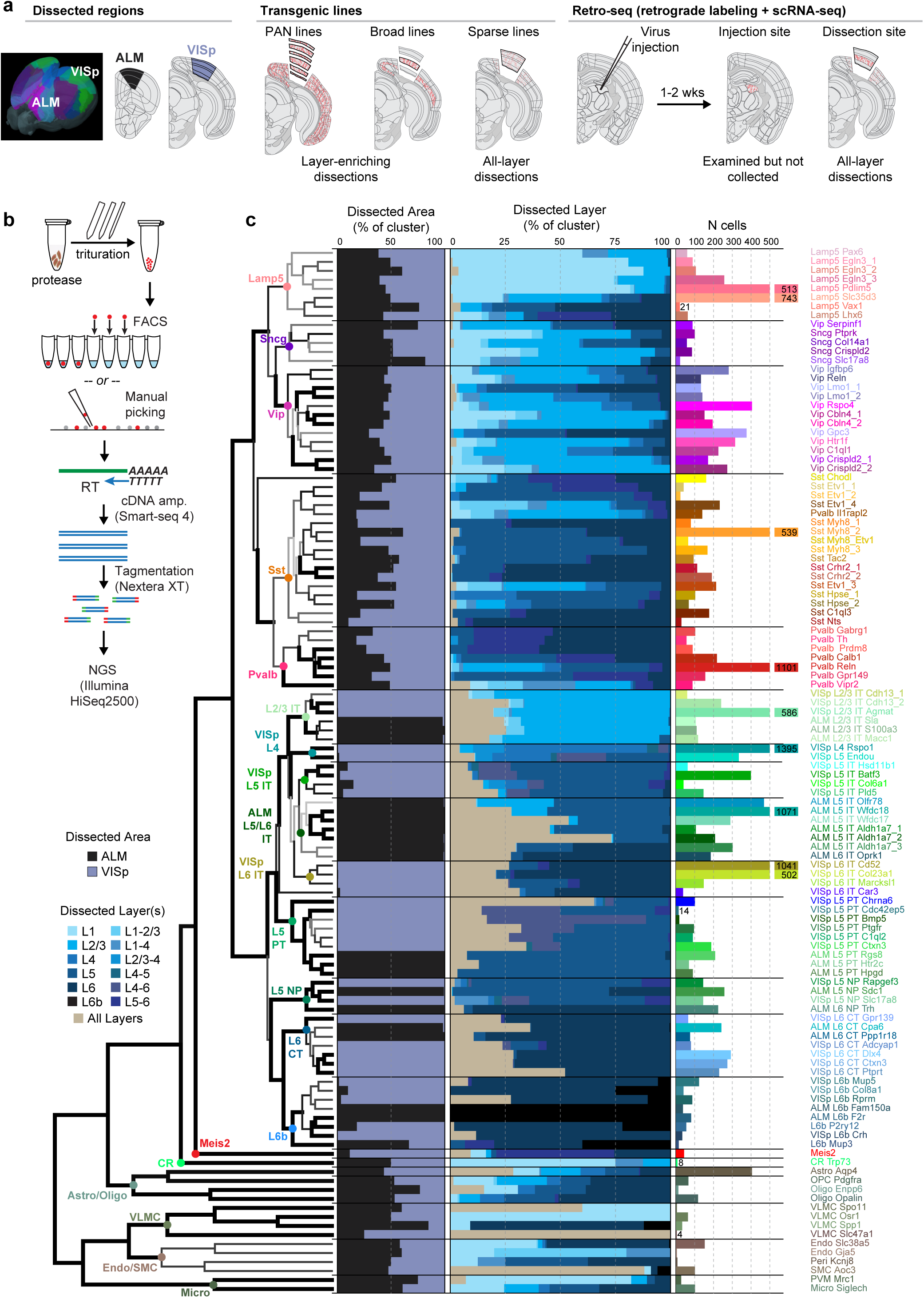
Cell type taxonomy in ALM and VISp cortical areas. (**a**) Cells were collected from the anterior lateral motor cortex (ALM) and primary visual cortex (VISp). For both areas, a combination of transgenically labeled cells and retrogradely labeled cells was used for scRNA-seq, in combination with layer-enriching or all-layer microdissections. (**b**) After dissection, samples were treated with a protease and triturated to generate a single-cell suspension. Individual cells were isolated by FACS or manual sorting, processed for reverse transcription (RT) and cDNA amplification (cDNA amp.) using SMART-Seq v4, followed by tagmentation and indexing. Indexed samples were then pooled for next-generation sequencing (NGS). (**c**) 50 GABAergic, 52 glutamatergic, and 14 non-neuronal types are organized into a hierarchical taxonomy on the basis of median cluster gene expression of 4,015 differentially expressed genes and 21,749 cells (12,714 from VISp, 9,035 from ALM). Hierarchical clustering was performed 100 times using multiscale bootstrap resampling to generate confidence scores for each branch point in the dendrogram (**Methods**). Dendrogram branches are colored based on their confidence scores (white = 0 to black = 1), and are thicker if the confidence score p-value < 0.05. Cell classes are labeled with colored dots at branch points in the dendrogram. The bar plots from left to right represent: the fraction of cells dissected from ALM (black) and VISp (bluegray); the fraction of cells from each of the layer-enriching dissections (lighter = upper layers, darker = lower layers; tan = all layers); and the number of cells contributing to each cell type in cluster-specific colors used throughout the paper.

A clear hierarchy of transcriptomic cell types and their relationships has emerged (**Fig. 1**). Consistent with previous reports (Tasic et al., 2016; Zeisel et al., 2015), the biggest differences in transcriptomic profiles are observed between non-neuronal and neuronal cells. We find that non-neuronal cells (n = 1,105 cells) are divided into two major branches according to their developmental origin: neuroectoderm-derived branch, which contains astrocytes and oligodendrocytes, and non-neuroectoderm-derived, which includes immune cells (microglia, perivascular macrophages), blood vessel-associated cells (smooth muscle cells, pericytes, and endothelial cells), and vascular leptomeningeal cells (VLMCs) (**Extended Data Fig. 9**). We identified previously described non-neuronal types, but also find pericytes and four different types of VLMCs with unique, previously unknown markers (**Extended Data Fig. 9**). The pericyte definition has been a matter of a debate due to their diverse morphologies and shared marker expression with smooth muscle cells, including *Acta2* (smooth muscle actin, α-SMA), *Pdgfrb*, and *Cpsg* (proteoglycan NG2) (Attwell et al., 2016). We find one transcriptomic cell type that specifically expresses *Kcnj8* and *Abcc9*, which have been identified as brain pericyte markers (Bondjers et al., 2006), and define additional markers uniquely expressed in this cell type (*Atp13a5*, *Art3*, *Pla1a*, and *Ace2*) that may help solidify pericyte identity in future studies (**Extended Data Fig. 9**).

Most neurons fall into two major branches corresponding to cortical glutamatergic and GABAergic types (**Fig. 1**). As exceptions, two very distinct types stand out: CR-Trp73, and Meis2, which are positioned in the taxonomy as two distant branches preceding the major glutamatergic/GABAergic split. Based on the marker expression and cell source, Meis2 type corresponds to the recently described *Meis2*-expressing GABAergic neuronal type of unknown function that is largely confined to the white matter, and originates from the embryonic pallial–subpallial boundary (Frazer et al., 2017). This type expresses the smallest number of genes (median = 3,863, **Extended Data Fig. 4b**) and is the only cortical GABAergic cell that reliably expresses the transcription factor *Meis2*. The other type corresponds to Cajal-Retzius (CR) cells based on their location in L1 and expression of *Trp73*, *Lhx5* and *Reln* (**Extended Data Fig. 6**), known markers of this cell type (Abellan et al., 2010; Kirischuk et al., 2014). Interestingly, CR cells of hippocampus and neocortex also originate from a distinct germinal zone, the cortical hem (Yoshida et al., 2006).

All other neuronal types fall into two major classes in accordance with the main neurotransmitter they express: GABA or glutamate. Every GABAergic neuronal type contains cells from both ALM and VISp (**Fig. 1**). That is, inhibitory GABAergic interneuron types are shared between these two cortical areas. In contrast, the glutamatergic types are segregated by region (with the exception of two L6b types and CR cells). Glutamatergic types segregate by global projections, cortical layer, and area, usually in that order (**Fig. 1**, see below).

### GABAergic cell type taxonomy

We define five subclasses within the GABAergic taxonomy: Lamp5, Sncg, Vip, Sst and Pvalb, and two distinct types Sst-Chodl, and Meis2 (**Fig. 1**). Within each subclass there are multiple types (**Fig. 2**). We represent the taxonomy derived from the current dataset by hierarchical clustering (**Fig. 1, Fig. 2c,d**) and constellation diagrams, which incorporate information on core and and intermediate cells (**Fig. 2a,b**). In addition, we illustrate layer-of-origin for cells collected from layer-enriching dissections (**Fig. 2c,d**), as well as the distribution of expression of individual marker genes within each type (**Fig. 2e,f**).

**Figure 2.**
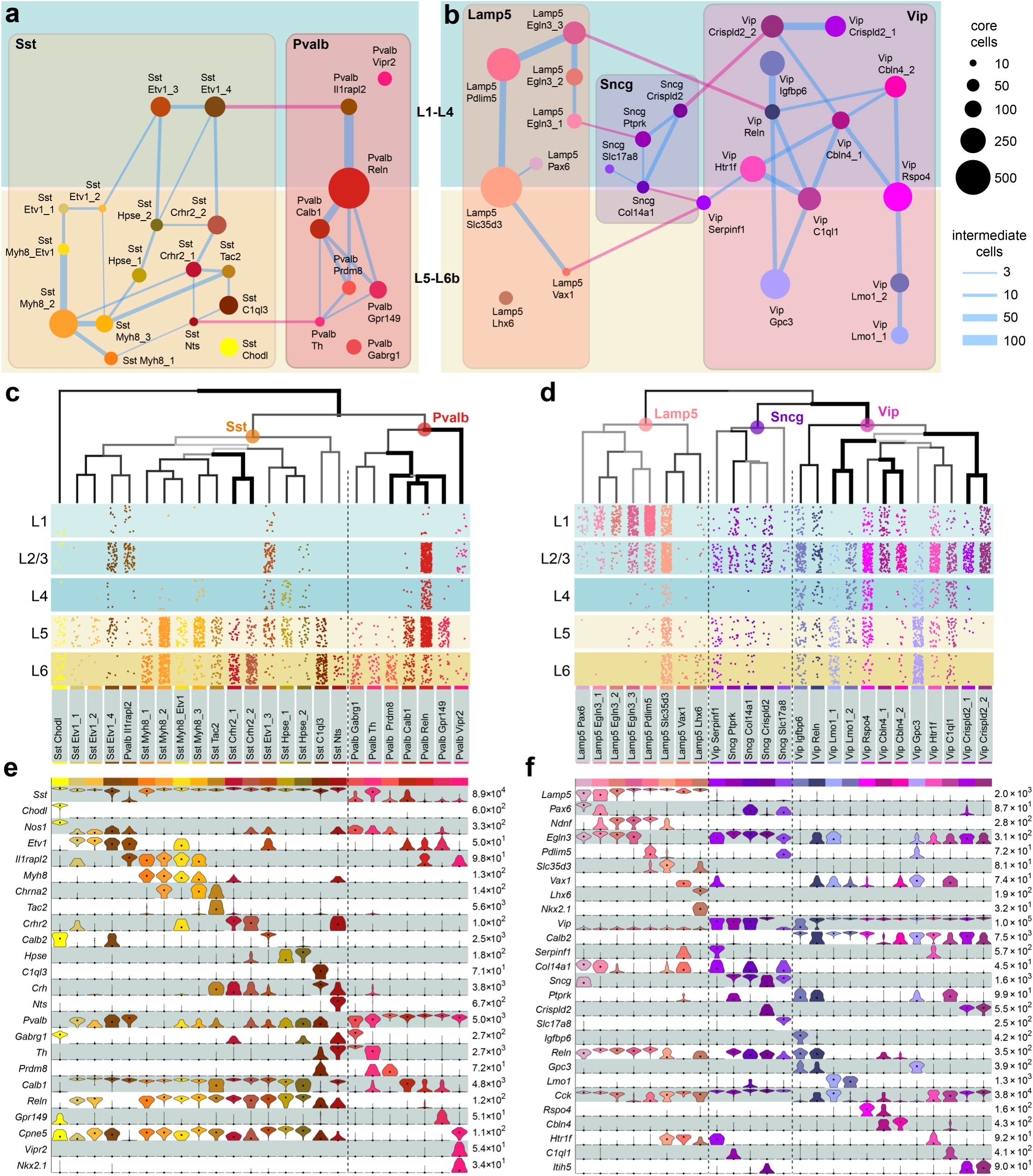
GABAergic cell types by scRNA-seq. (**a,b**) Constellation diagrams for Sst/Pvalb and Lamp5/Sncg/Vip types. Core cells (N = 8,040) are represented by discs, and intermediate cells (N = 1,259) by lines. Intermediate cells connecting different classes (e.g., Sst and Pvalb) are labeled pink. No intermediate cells exist between Sst/Pvalb and Lamp5/Sncg/Vip classes. (**c,d**) Dendrograms represent overall gene expression similarity between types based on median gene expression in each cell type for differentially expressed marker genes (as in **Fig. 1**). Below the dendrograms, spatial distribution of each type was inferred from layer-enriching dissections: each dot represents a single cell from a layer-enriching dissection, dots are positioned at random within each layer. (**e,f**) Marker gene expression distributions in single cells within each cluster are represented by violin plots. Each row represents a single gene, and values within rows are normalized between 0 and the maximum expression value for each gene (shown at the right edge of each row) and displayed on a log_10_ scale. Median values are shown as black points within each violin.

The GABAergic cells are divided into two major classes, mostly corresponding to their developmental origin in the medial ganglionic eminence (MGE: Pvalb and Sst types) or caudal ganglionic eminence (CGE: Lamp5, Sncg and Vip types). The developmental origins of some of the types have not been established; for example, Lamp5-Lhx6 type may originate from the MGE (see Discussion). Many inhibitory types display layer-specific enrichments (**Fig. 2c,d**). The Sst and Pvalb subclasses within the Sst/Pvalb constellation are connected at two different points: one connection occurs between two upper-layer types (Sst-Etv1-4 and Pvalb-Ilrapl2) and the other between two deeper-layer types (Sst-Nts and Pvalb-Th) (**Fig. 2a**). The CGE-derived interneurons, united under the umbrella of *Htr3a-GFP* transgene expression (Rudy et al., 2011), fall into Lamp5, Vip, and Sncg subclasses that are represented by three interconnected neighborhoods in the constellation diagram (**Fig. 2b**). The complicated landscapes are the result of many genes expressed in a combinatorial and graded fashion (Extended Data Figs. 5 and 6), resulting in high co-clustering frequencies (**Extended Data Fig. 3a)** and many intermediate cells.

The GABAergic transcriptomic taxonomy overlaps with previously reported interneuron types based on marker gene expression, transgenic lines, published Patch-seq and other scRNA-seq data (**Extended Data Table 2, Extended Data Figs. 10 and 11**). The transcriptomic taxonomy now provides a global context for these more focused studies of interneuron types. Some of the most distinct types are Meis2 (discussed above), and Sst-Chodl, which corresponds to *Nos1*-expressing long-range projecting interneurons, based on marker gene expression, location, Cre line labeling, and other RNA-seq data (**Extended Data Table 2, Extended Data Figs. 10a, 11**) (He et al., 2016; Paul et al., 2017; Tasic et al., 2016).

For the Pvalb subclass, we confirm that the Pvalb-Vipr2 type (named Pvalb-Cpne5 in our previous study (Tasic et al., 2016)), corresponds to chandelier cells by high confidence mapping of the recently reported chandelier cell 1 (CHC1) type RNA-seq data (Paul et al., 2017) to our Pvalb-Vipr2 type (**Extended Data Fig. 10a**). We also used the newly discovered genetic marker *Vipr2* to develop *Vipr2-IRES2-Cre* for access to chandelier cells without requiring tamoxifen induction, which is necessary for chandelier cell labeling by the *Nkx2.1-CreERT2* line (**Extended Data Figs. 11, 12a-f**). Most other Pvalb types correspond to basket cells (**Extended Data Fig. 10a,b)** (Paul et al., 2017), except for two L6-enriched types (Pvalb-Th and Pvalb-Gabrg1) whose identities are still elusive.

Within the Lamp5/Vip/Sncg subclasses, we find evidence for correspondences to neurogliaform, bipolar, single bouquet, and CCK basket cells (**Extended Data Table 2**). The newly defined Sncg subclass corresponds to the *Vip*+ and *Cck*+ multipolar or basket cells and is distinct from the cells of the Vip subclass that are also *Calb2*+ and have mostly bipolar morphologies (**Fig. 2f, Extended Data Fig. 10a**)(He et al., 2016; Paul et al., 2017; Rudy et al., 2011). We previously assigned neurogliaform cell (NGC) identity to Ndnf types (Tasic et al., 2016), which correspond to several current Lamp5 types (**Extended Data Fig. 7**). We confirm this finding by computational mapping of Patch-seq data (Cadwell et al., 2016) to our data (**Extended Data Fig. 10d-f)** to find excellent correspondence of NGCs to our Lamp5-Pdlim5 and Lamp5-Slc35d3 types. In addition, we find that single bouquet cells (SBCs) map to the Lamp5-Egln3_2 type (**Extended Data Fig. 10d,e**), and find a possible transitional SBC/NGC type, Lamp5-Egln3_3 (**Extended Data Fig. 10d**).

The Lamp5-Lhx6 type is unusual because it clusters with other Lamp5 types, but is the only one expressing transcripts encoding the typical MGE transcription factors, *Nkx2.1* and *Lhx6.* Consistent with the marker gene expression, this type is labeled by *Nkx2.1-CreERT2* (**Extended Data Fig. 11**) and was isolated by Paul *et al.* from the same Cre line (**Extended Data Fig. 10a-c**). Intriguingly, we find that the chandelier type 2 cells (CHC2) RNA-seq data (Paul et al., 2017) map exceptionally well to our Lamp5-Lhx6 type (**Extended Data Fig. 10a,b**), which is transcriptomically most related to Lamp5 neurogliaform types.

The Sst subclass exhibits diverse transcriptomic types, currently with only a few clear correspondences with known *Sst*+ morphological types, in addition to the above-mentioned Sst-Chodl type. The Sst-Etv1_3 type corresponds to the *Sst*+ and *Calb2*+ L2/3 Martinotti cells (**Fig. 2e, Extended Data Fig. 10a)** (He et al., 2016; Paul et al., 2017; Rudy et al., 2011). Some of the Sst-Myh8 types (Sst-Myh8_2 and Sst_Myh8_3) and the Sst-Tac2 type express the *Chrna2* marker gene, and thereby likely correspond to L5 Martinotti cells (Hilscher et al., 2017).

### Glutamatergic cell type taxonomy based on scRNA-seq and projections

Most excitatory neurons in the cortex project to other brain regions, including other cortical regions, and genetic markers have been correlated with projectional properties (Molyneaux et al., 2007; Sorensen et al., 2015) (Harris and Shepherd, 2015). To inquire how transcriptomic and projectional identities of excitatory neurons correlate with each other, we included 2,677 cells labeled by retrograde injections (Retro-seq dataset, **Fig. 3a**) in our single cell transcriptomics dataset. The projection targets of VISp and ALM (**Fig. 3b, Extended Data Fig. 13)** were selected based on the Allen Mouse Brain Connectivity Atlas (Oh et al., 2014) and other anatomical data (Li et al., 2015). Retrogradely labeled cells were isolated after viral infections, and were processed for single cell RNA-seq through the same experimental and data processing pipeline, including joint clustering with cells isolated from transgenic lines. The clustering was performed blind to the identity of retrogradely labeled cells, and these cells were only used post-clustering to represent the correspondence between projectional and transcriptomic properties.

**Figure 3.**
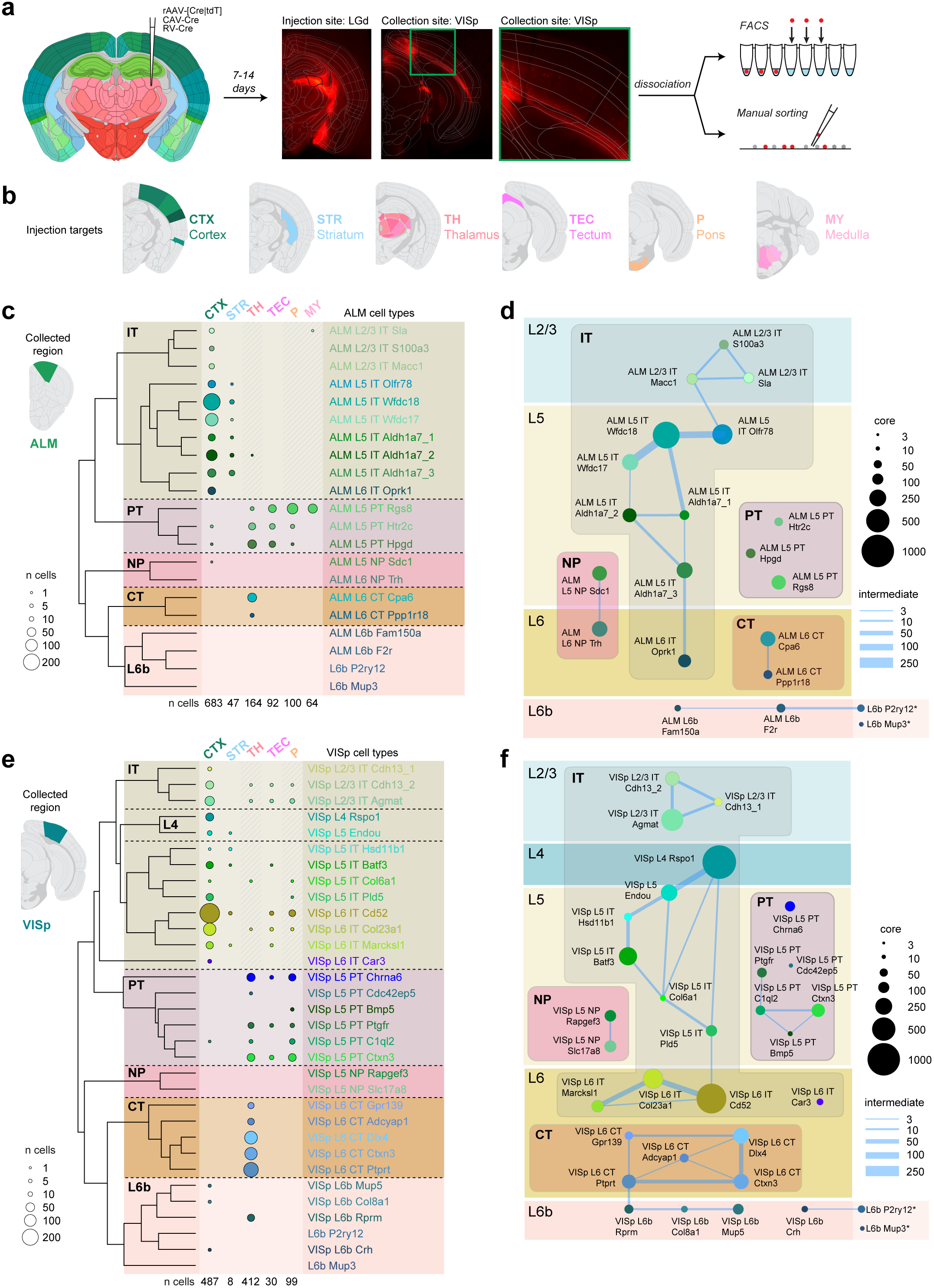
Glutamatergic cell types by scRNA-seq and projections. (**a**) For each retrograde tracing experiment, a virus was injected into a target site. 7-14 days after injection, brains were sectioned, and the injection site was imaged to determine injection specificity. Cells were collected from the collection site (ALM or VISp) by microdissection, tissue was dissociated and single cells sorted by FACS or manually. (**b**) Injection targets are grouped into target regions: Cortex (CTX), Striatum (STR), Thalamus (TH), Tectum (TEC), Pons (P), or Medulla (MY). More details about injections are available in **Extended Data Fig. 13. (c)** Dendrogram representing all excitatory cell types in ALM followed by numbers of cells (represented by discs) originating from retrograde labeling by injections into regions labeled on top. Shaded regions denote cells that may have been labeled unintentionally, directly or retrogradely, through the needle (injection) tract. (**d**) Constellation diagram for all cells from ALM glutamatergic cell types. Disc area represents the total number of core cells assigned to each clusters, and edge weights represent the number of intermediate cells. (**e**) Same as (c), but for VISp. VISp cells were not collected from injections into medulla (MY). (**f**) Same as (d), but for VISp. *, L6b-P2ry12 and L6b-Mup3 types are found in both ALM and VISp.

The presence of retrogradely labeled cells in transcriptomic clusters revealed their projection patterns (**Figs. 3 and 4**). Intratelencephalic (IT) types, constituted the largest branch, whereas pyramidal tract (PT) types, near-projecting (NP) types, and corticothalamic (CT) types constituted separate branches. All branches contained related types from both areas (**Fig. 4a**). The only subclass dominated by layer identity was L6b subclass.

**Figure 4.**
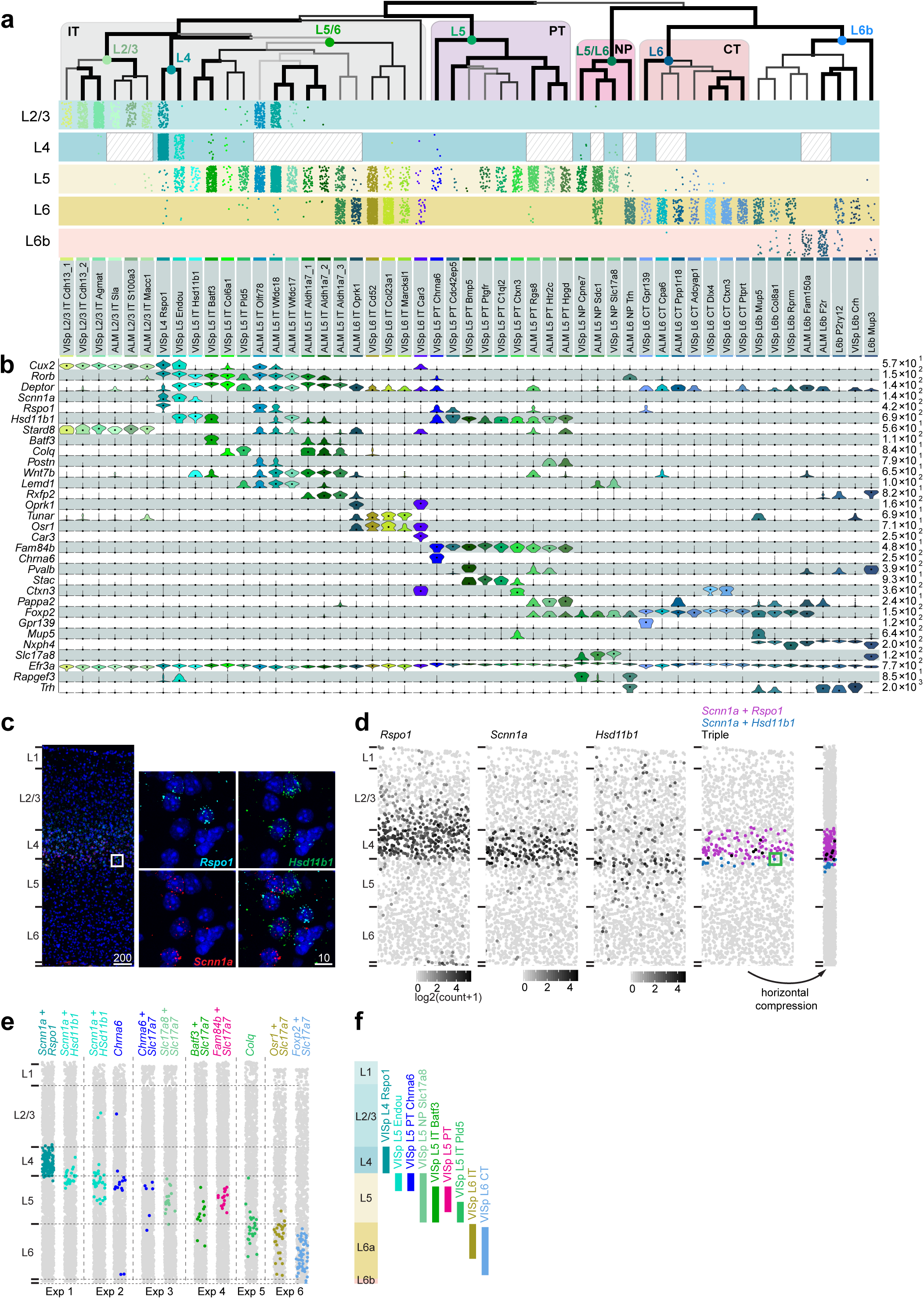
Glutamatergic cell types and markers. (**a**) A dendrogram representing taxonomy of 51 glutamatergic clusters based on 11,377 total cells and median gene expression of 1,289 genes per cluster. Below, cells from layer-enriching dissections that belong to these types are represented as dots (randomly within each layer). ALM does not contain L4 (indicated by hashed rectangles). (**b**) Violin plots represent distributions of individual marker gene expression in single cells within each cluster. Rows are genes, median values are black dots, and values within rows are normalized between 0 and the maximum expression value for each gene (right edge of each row) and displayed on a log_10_ scale. (**c**) Single molecule fluorescence RNA in situ hybridization with RNAscope to validate marker expression and cell type distribution. Example image shows fluorescent spots that correspond to *Rspo1*, *Hsd11b1*, and *Scnn1a* mRNA molecules in a 10-μm coronal VISp section. Scale bars are in μm. (**d**) Example of processed RNAscope data obtained by generating a maximum projection of a montage of confocal z-stacks, identifying nuclei, quantifying the number of fluorescent spots, and assigning spots to each nucleus by CellProfiler (Lamprecht et al., 2007). Each dot in the panel represents a cell in the same VISp region as in **c**, plotted according to the detected nucleus position. The first three panels show cells shaded according to the quantified number of spots per cell: *Rspo1* and *Scnn1a* mRNAs are enriched in L4 and *Hsd11b1* at the L4/5 border. The fourth panel shows location of cells co-labeled with two or more probes and confirms scRNA-seq data: coexpression of *Rspo1* and *Scnn1a* is expected in VISp L4-Rspo1 type and coexpression of *Hsd11b1* and *Scnn1a* in VISp L5-Endou type (see **a,b**). The green square in (**d**) corresponds to white square in (**c**). The fifth panel in (**d**) is a condensed plot created to emphasize layer enrichment of examined cells. (**e**) Condensed plots for RNA scope experiments (Exp) in VISp for select excitatory cell type markers. Layers were delineated based on cell density. (**f**) A schematic of laminar distributions of VISp glutamatergic types according to experiments in (**e**).

IT types from both areas are present in all layers except L1, they express previously reported layer-specific or enriched markers (**Fig. 4a,b, Extended Data Fig. 6**) and map well to our previously detected types (**Extended Data Table 2, Extended Data Fig. 7**) (Tasic et al., 2016). They display a significant number of intermediate cells, which connect types within a layer or in neighboring layers (**Fig. 3d,f**). We also define a distinct IT type, VISp L6 IT-Car3, which expresses a unique combination of many markers including *Car3*, *Oprk1* and *Nr2f2* (**Fig. 4a,b**), some of which have been previously detected in the claustrum (Wang et al., 2017). Examination of the Allen Brain Atlas RNA ISH data shows that these genes are expressed in a sparse population of L6 neurons located in the lateral portion of VISp and continuing laterally towards the claustrum (**Extended Data Fig. 14**).

We performed anterograde tracing by Cre-dependent adeno-associated virus (AAV) in select Cre lines (**Fig. 5a)** to examine the cortico-cortical projection patterns of different IT types in more detail. We find cases where different types project to different cortical areas, or different types project to different layers within the same projection area. L5 and L2/3 IT types as labeled in *Tlx3-Cre*_*PL56*, and *Cux2-IRES-Cre* lines (**Extended Data Fig. 11)** display extensive long-range projections that cover all layers with preference for upper layers (**Fig. 5b,c**). In contract, L6 IT types, labeled in the newly generated *Penk-IRES2-Cre-neo* line (**Extended Data Fig. 11**), project to many of the same areas as the L2/3 and L5 IT types, but their projections are confined to lower layers in all areas examined, including higher visual areas and contralateral VISp and ALM (**Fig. 5d**). As revealed by our Retro-seq data (**Extended Data Fig. 13b**), we confirm that L4-Rspo1 type, which represents most cells labeled by *Nr5a1-Cre* (**Extended Data Fig. 11)** projects to contralateral VISp (**Fig. 5e**). Interestingly, this projection is observed only when injection is performed in the most anterior portion of VISp (compare left and right panels in **Fig. 5e**).

**Figure 5.**
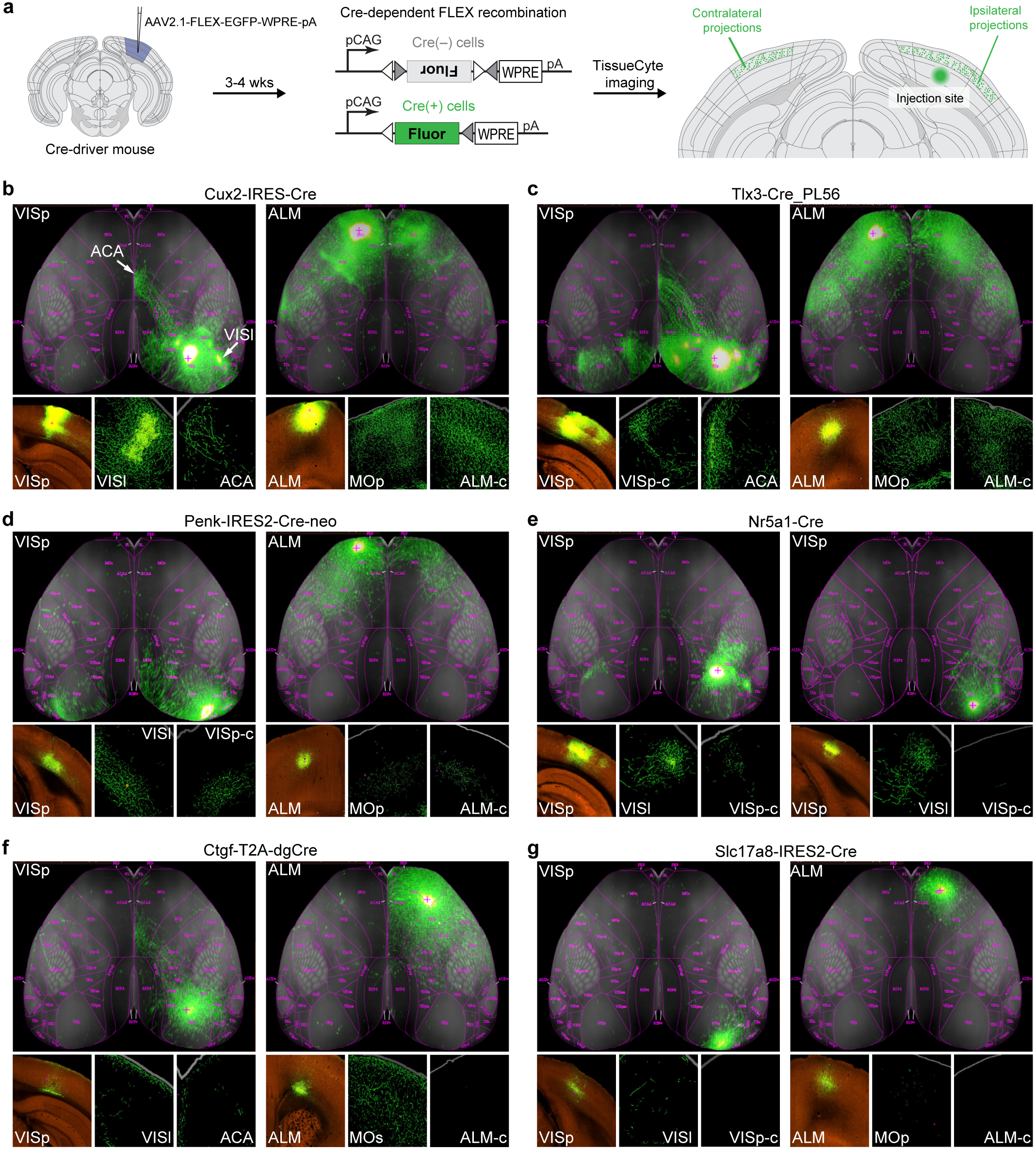
Anterograde tracing reveals area- and layer-specific cortico-cortical projections of transcriptomic types. (**a**) Anterograde tracing workflow: Cre-dependent adeno-associated virus (AAV) was injected into VISp or ALM in different Cre-driver mice. 3 weeks post-injection, the brains were imaged by the TissueCyte 1000 system, the image datasets were registered to Allen Mouse Common Coordinate Framework (CCF), and the GFP fluorescent signals were segmented to reveal GFP-labeled axon fibers throughout the entire brain (Oh et al., 2014). (**b-g**) Left and right panels represent injections into VISp and ALM, respectively, for (**b**) *Cux2-IRES-Cre*; (**c**) *Tlx3-Cre*_*PL56*; (**d**) *Penk-IRES2-Cre-neo*; (**f**) *Ctgf-T2A-dgCre;* and (**g**) *Slc17a8-IRES2-Cre* animals. (**e**) Left and right panels represent two different injection locations within VISp for *Nr5a1-Cre* animals. Top image is top-down view of segmented GFP signals throughout the cortex, bottom-left image is a raw fluorescent image of the injection site; middle and right images are segmented GFP signals from representative projection target areas in contralateral (labeled with ‘c’) or ipsilateral side. VISl, lateral visual area; VISp-c, contralateral VISp; ALM-c, contralateral ALM; ACA, anterior cingulate area; MOp, primary motor area; MOs, secondary motor area.

Cortico-subcortical projecting, or pyramidal-tract (PT) neurons, which are the descending output neurons in L5, share a separate branch in the taxonomy tree and express previously known markers like *Bcl6* (Sorensen et al., 2015) and a new marker *Fam84b* (**Fig. 4a,b**). The PT transcriptomic types in ALM correspond to two projection classes (Economo et al., 2017, accompanying paper): two transcriptomic types project to thalamus (ALM L5 PT-Htr3c and -Hpgd), whereas the third projects to the medulla (ALM L5 PT-Rgs8, **Extended Data Fig. 13a**). These PT types have distinct functions in planning and initiating voluntary movements (Economo et al., 2017, accompanying paper). Likewise, it appears that PT types from VISp display differential subcortical projections (**Extended Data Fig. 13b**); however new tools specific for individual L5 PT types will be needed to confirm this possibility.

Cortico-thalamic (CT) L6a types share the transcription factor marker *Foxp2*, and may have type-specific preferences for different thalamic nuclei: VISp L6 CT-Dlx4 and -Ctxn3 types appear to more preferentially project to LGd, whereas VISp L6 CT-Ptprt roughly equally targets LP, LD and LGd (**Extended Data Fig. 13b**).

L6b types share marker genes for subplate neurons, such as *Cplx3, Ctgf* and *Nxph4* (Ayoub and Kostovic, 2009; Lein et al., 2007; Zeng et al., 2012). We find two L6b types that are shared between the two areas: L6b-Mup3 and L6b-P2ry12. Based on the Retro-seq dataset, one of the L6b types (VISp L6b-Rprm), which shares a number of markers with L6a CT types including *Rprm* and *Crym*, also projects to thalamus. The dendrograms (**Figs. 1c and 3e**) capture only the dominant relationship between L6b types, whereas the constellation diagram can accommodate more complicated relationships showing intermediate cells between VISp L6b-Rprm and VISp L6 CT-Ptprt (**Fig. 3f**). Anterograde tracing in *Ctgf-T2A-dgCre* mice shows that L6b neurons have predominant projections into L1 that spread to many neighboring cortical areas in both VISp and ALM (**Fig. 5f**). In agreement with the Retro-seq data (**Extended Data Fig. 13b**), we also find sparse long-range projections from anterior VISp to anterior cingulate area (ACA, **Fig. 5f**, left panel).

We define four related types (two per area) in L5/6 that express distinct markers including *Tshz2*, *Slc17a8* and *Rapgef3*. Based on the Retro-seq dataset, they do not project to any of the assayed areas (**Fig. 3c,e**). To evaluate this finding by anterograde tracing we used a newly generated Cre line, *Slc17a8-IRES2-Cre*. Anterograde tracing of neurons labeled by this line revealed no long-range projections, except for a few axons in neighboring areas (**Fig. 5g**). Therefore, we name this subclass “near-projecting” (NP). Some of these cells likely correspond to previously reported *Slc17a8*+ L5 cells, whose projections could not be established by retrograde labeling (Sorensen et al., 2015), as well as cells labeled by *Efr3a-Cre_NO108* (Kim et al., 2015). Although *Efr3a* mRNA is ubiquitously detected in all neurons, its expression is about 5-fold higher in NP types compared to others (**Fig. 4b);** this may contribute to preferential labeling of the NP types by this BAC transgenic Cre line.

We confirmed the spatial locations of the transcriptomic types in VISp by single-molecule FISH with class-or type-specific marker genes as probes (**Fig. 4c-f**). Our result (**Fig. 4-f**) corroborates previous evidence showing that L5 IT and L5 PT cells are not clearly separated into 5a and 5b sublayers in the visual cortex in contrast to their sublayer-specific segregation in the primary somatosensory cortex (Groh et al., 2010).

### Cell types: area-specific divergences

To assess differences in the number of differentially expressed genes between most closely-related types, we used the best match between types for excitatory types (**Fig. 6a,b)** and, as a comparison, we split the inhibitory clusters into their area-specific (ALM- and VISp-) portions. We then performed differential gene expression search to count the number of differentially expressed genes between these best-matched pairs. We find that inhibitory neurons from the two areas belonging to the same cluster have at most 12 differentially expressed genes, while the best matched excitatory types between VISp and ALM have a median of 100 differentially expressed genes (**Fig. 6a, Extended Data Table 3**).

The method of defining the best match among excitatory types between the two areas (**Fig. 6b**), revealed additional patterns. Regardless of the direction of mapping (VISp cells onto ALM types or vice versa), very few one-to-one correspondences between types were observed. Several-to-several mappings dominate, but are almost completely confined to each major branch (subclass), confirming similarity at higher hierarchical level between ALM and VISp types. We identify a large set of genes with differential enrichment in these two areas, further highlighting substantial inter-areal differences in excitatory cell populations (**Fig. 6c**). Interestingly, we find more ALM-enriched genes. We confirm the area-specific expression of several genes by RNA ISH from the Allen Brain Atlas (**Fig. 6e**).

**Figure 6.**
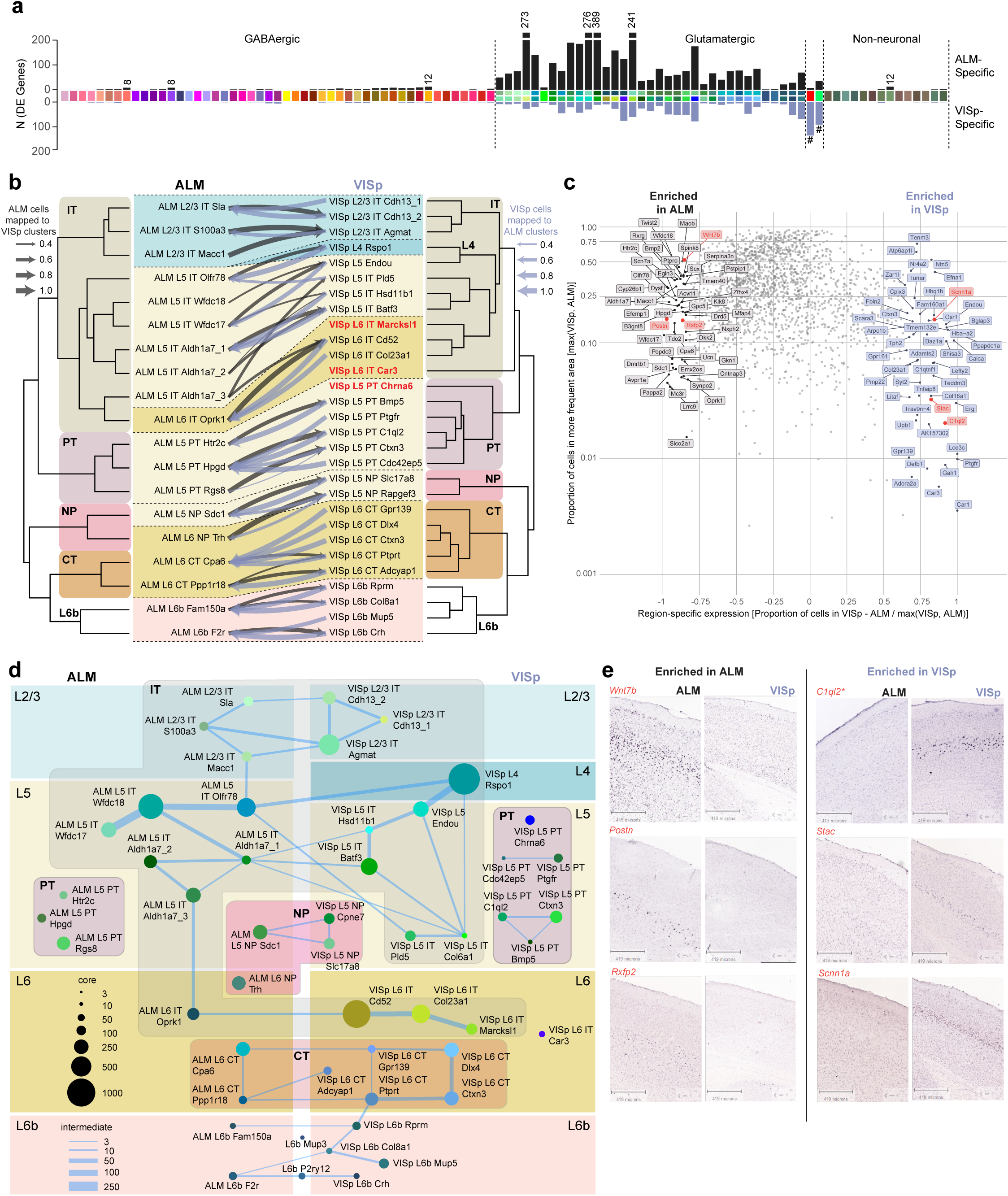
Comparison of gene expression differences among types across cortical areas. (**a**) Number of differentially expressed genes between best matching cell types (for types that are different between areas) or ALM- and VISp-portions of the same type for shared types. (**b**) Mapping glutamatergic cells from ALM onto VISp glutamatergic cell types (gray arrows) using a random forest classifier trained on VISp types, and vice versa (blue-gray arrows; **Methods**). The fraction of cells that mapped with high confidence onto clusters from the other region is represented by the weight of the arrows. (**c**) Genes that are specific to ALM (gray) or VISp (blue-gray) based on the proportion of cells in each region that express each gene. Two measures are shown here: a ratio of proportions (proportion of cells in ALM – proportion in VISp divided by whichever is higher, x-axis, **Methods**) and the proportion of cells in whichever region has a greater proportion of cells expressing each gene (y-axis). (**d**) Joint constellation diagram for ALM and VISp. (**e**) RNA ISH from the Allen Mouse Brain Atlas for select genes (red in **b**) confirms areal gene expression specificity.

We also construct a constellation diagram for excitatory types from both areas (**Fig. 6d**), revealing similar correspondences to the mapping in **Fig. 6b**: correspondence is based both on layer and projectional type, and there are no intermediate cells that connect types from non-adjacent layers.

Inhibitory neurons belonging to the same cluster and segregated by area display at most 12 differentially expressed genes (**Fig. 6a**). Using *t*-Distributed Stochastic Neighbor Embedding (t-SNE), we observe more pronounced segregation of cells by region for Sst/Pvalb types compared to Vip/Lamp5 types (**Fig. 7a,b**), with both being substantially lower than segregation observed for glutamatergic types (**Fig. 7c**). An example of regional expression differences within the Sst Etv1_3 and Sst_Crhr2_1 clusters (**Fig. 7d**) was observed for *Crh* and *Calb2* genes. These genes are co-expressed in some cells in Sst Etv1_3 type only in ALM, and in Sst Crhr2_1 type only in VISp (**Fig. 7e)**. We examined this finding by single-molecule RNA FISH using probes against *Sst*, *Calb2*, and *Crh* (**Fig. 7f,g**). As predicted by cell type identity and cell type distribution, we find triple positive cells by RNA FISH in ALM only in upper layers, likely belonging to Sst Etv1_3 type (**Fig. 7e,f**). In contrast, the triple positive cells in VISp are only found in lower layers, likely corresponding to Sst Crhr2_1 (**Fig. 7e,g)**.

**Figure 7.**
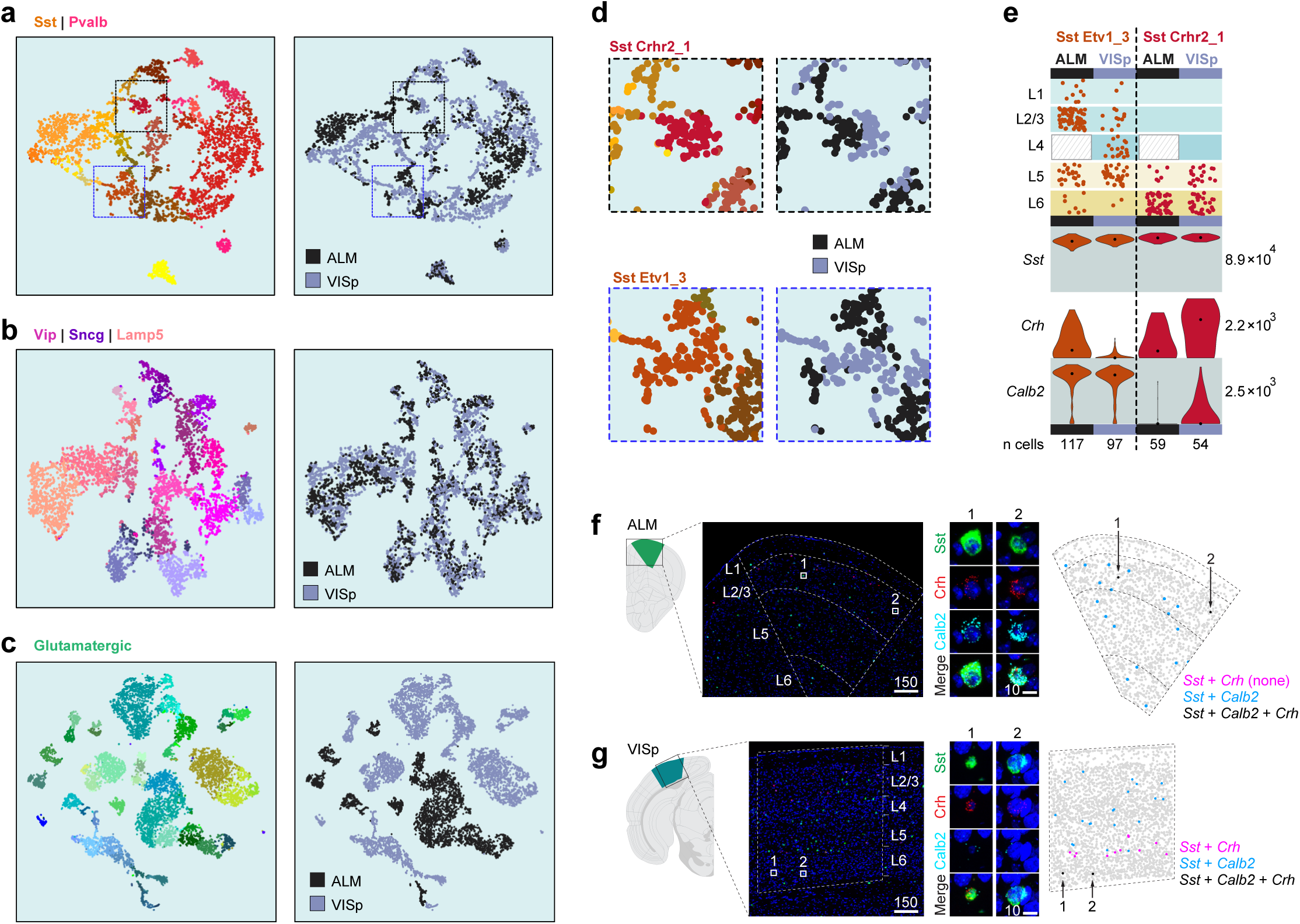
Comparison of gene expression differences within inhibitory types across cortical areas. (**a-c**) *t*-SNE plots for Sst/Pvalb (4,259 cells, 747 differentially expressed genes), Lamp5/Sncg/Vip (5,006 cells, 756 differentially expressed genes) and excitatory types (11,337 cells, 1,289 differentially expressed genes) showing cells labeled in cluster colors on the left and area-of-origin on the right. Glutamatergic cells show most dramatic segregation, whereas Sst/Pvalb types show small but noticeable segregation within the same type based on the area. This is less obvious for the Lamp5/Sncg/Vip types. (**d**) Areas from the Sst/Pvalb *t*-SNE plots were enlarged to show partial segregation of cells within two Sst types by area of origin. (**e**) Differential gene expression in the same transcriptomic type divided by area confirmed by smFISH (**f,g**).

## DISCUSSION

We use single-cell transcriptomics to uncover principles of cell type diversity in the neocortex and their variation across functionally distinct areas. By employing a variety of recombinase driver lines, we achieved both broad coverage and enrichment for rare or fragile cell types to define a near-complete repertoire of transcriptomic cell types in two functionally distinct cortical areas, VISp and ALM. By combining retrograde labeling from multiple VISp and ALM output areas, we defined correspondence between transcriptomic identity and neuronal projection properties. The transcriptomic types were organized in a hierarchical taxonomy through unsupervised clustering analyses, and fell into major branches (cell classes) and terminal leaves (cell types) that are either novel or correspond to known cell classes/types. Consequently, this dataset provides comprehensive gene expression profiles and genetic markers for these novel and existing cell classes/types that will enable genetic access to them for future studies from molecular to functional levels.

We define 116 transcriptomic types, 86 types in ALM and 97 in VISP, 67 of which are shared between the two areas. The shared types are non-neuronal and inhibitory neuronal types, whereas most excitatory types are area-specific (exceptions are the CR cells and two L6b types). Although previous studies reported similarities of neocortical and hippocampal interneurons, as well as non-neuronal cells (Zeisel et al., 2015), we report this finding with a more sensitive technique including substantially higher gene detection. In fact, we find statistically significant gene expression differences between ALM and VISp portions of many interneuron clusters (**Figs. 6a and 7**); however these differences are still insufficient to cause cluster separation into VISp- and ALM-specific interneuron subtypes with our statistical criteria (Methods). In contrast, we readily detect a large number of genes that distinguish each pair of matched excitatory types across these two cortical areas (**Fig. 6a)**.

This dichotomy correlates well with area-specific connectivity patterns as well as the cells’ developmental origins. Most excitatory neuronal types in VISp or ALM project to different cortical and subcortical target areas (**Fig. 5, Extended Data Fig. 13)**, forming unique circuits involved in distinct neural coding processes. In contrast, nearly all inhibitory interneurons form local connections with nearby excitatory or inhibitory neurons, and play mostly modulatory functions within the local circuits. Although the inhibitory types are shared between areas, their area-specific functions may be achieved through the variations in cell type abundances across areas (Kim et al., 2017).

The developmental origins of these different neuronal types correlate with their uniqueness or commonality. It has been shown that most excitatory neurons are born locally within the ventricular/subventricular zone (Gao et al., 2014) that is pre-patterned with developmental gradients (O’Leary et al., 2007) and further segregated into areas through differential thalamic input early in development (Chou et al., 2013; Moreno-Juan et al., 2017; Vue et al., 2013). In contrast, GABAergic interneurons are MGE- and CGE-derived, Meis2 interneurons originate from the palial-subpalial boundary (Frazer et al., 2017), and Cajal-Retzius cells originate from the cortical hem (Yoshida et al., 2006). Similarly, we wonder if the two L6b types that are shared across areas correspond to a subset of subplate neurons that originate from the rostro-medial telencephalic wall, which is a known embryonic source for a subset of subplate neurons that are distinct from other subplate neurons generated within the local ventricular/subventricular zone (Hoerder-Suabedissen and Molnar, 2015; Pedraza et al., 2014).

We observe both discontinuous and continuous gene expression variation among and within cell types. Continuous variation poses a challenge to cell type categorization, as clustering methods are designed to rely on discontinuous variation. We therefore used post-clustering classifiers and constellation diagrams to describe both types of variation within the cell type landscape. These landscapes may evolve as the perceived vs. actual discreteness (or discreteness vs. continuity) in cell type definition depend on gene detection, cell sampling and noise estimates (Tasic et al., 2017). For example, in this study, we detect a continuum of phenotypes within the L4-Rspo1 type that corresponds to three highly related previous types (L4 Ctxn3, L4 Scnn1a, and L4 Arf5), which were connected by many intermediate cells (**Extended Data Table 2, Extended Data Fig. 15**) (Tasic et al., 2016). Therefore, heterogeneity is still present, but its description changed from discrete to continuous.

The hierarchical organization of cell types captures the adult similarity in gene expression with high-level branches representing major, distinct cell classes and finer branches representing more closely related types. In many cases, it also represents developmental, and potentially evolutionary ontology of cell types. However, this does not necessarily have to be the case for every type – developmental origins need to be experimentally established. For example, we find that *Nkx2.1-CreERT2* labels not only chandelier cells but also a type that was presumed to be a second chandelier type – CHC2 (Paul et al., 2017). However, we find that based on its transcriptomic signatures, CHC2 is likely a neurogliaform type (Lamp5-Lxh6). The embryonic origin of this potential MGE-derived type would still have to be tested by lineage tracing, but it is not unreasonable to consider that the neurogliaform phenotype could arise through two different developmental pathways in an example of developmental convergence. For example, in *C. elegans*, morphologically and connectionally highly similar bilateral neuronal pairs are derived from different lineages (Hobert, 2014).

Our current hierarchical cortical cell type taxonomy is largely based on transcriptomic information that is overlaid with projectional properties. In the future, it will be important to correlate transcriptomic information with morphological, physiological, connectional, and functional properties of individual cells to define which transcriptomic features correspond to other phenotypes. As a particularly compelling example, we show that L5 pyramidal tract (PT) transcriptomic clusters correspond to neuronal types with distinct projection patterns and different functions (Economo et al., 2017, accompanying paper). We expect that as new data become available, the cell type taxonomy will be further refined. This dataset provides a foundation for understanding cortical cell type diversity and dissecting circuit function.

## ACKNOWLEDGEMENTS

We would like to thank M. Chillon Rodrigues (Universitat Autònoma de Barcelona) for providing CAV-Cre, Alla Karpova (Janelia) for providing AAV2-retro, Ali Williford for technical assistance, and the Department of In vivo Sciences for mouse husbandry. This work was funded by the Allen Institute for Brain Science, and by US National Institutes of Health grants R01EY023173 and U01MH105982 to H.Z. The authors thank the Allen Institute founder, Paul G. Allen, for his vision, encouragement and support.

## AUTHOR CONTRIBUTIONS

H. Z. and K.S. conceptualized the VISp and ALM cell type comparative study. H.Z. and B.T. designed and supervised the study. K.S. defined ALM coordinates based on loss-of function experiments. K.A.S. organized and managed single cell RNA-seq pipeline. D.B., J.G., K.L., C.R, M.T. and T.K.K. performed single-cell RNA-seq. Z.Y., L.T.G. and B.T. performed transcriptome data analysis with contributions from O.F., O.P., T.B., V.M., J.M., A.S. and M.H. I.W. and A.C. provided retrograde tracing viral vectors. J.A.H., T.N.N., K.E.H. and P.G. conducted retrograde and anterograde viral tracing experiments. B.P.L., N.D., T.C., S.P., E.B., M.K., N.V.S. and D.H. performed single-cell isolation. T.N.N. and E.G. performed RNA *in situ* hybridization with RNAscope. L.M. and T.L.D. generated transgenic mice. J. Pendergraft provided animal genotyping and R.L. provided mouse colony management. A.B. and J. Phillips managed pipeline establishment. K.S., M.E., S.V. and L.L. provided retrogradely labeled and manually collected single cells from ALM. S.M.S. provided program management support. H.Z. and E.L. led the Cell Types Program at the Allen Institute. C.K. and A.J. provided funding, institutional support and management. L.G., Z.Y., T.N.N., and B.T. prepared the figures. B.T. and H.Z. wrote the manuscript with contributions from C.K., K.S., L.T.G., T.N.N., and Z.Y., and in consultation with all authors.

## METHODS

### Mouse breeding and husbandry

All procedures were carried out in accordance with Institutional Animal Care and Use Committee protocols 1508, 1510, and 1511 at the Allen Institute for Brain Science. Animals were provided food and water *ad libitum* and were maintained on a regular 12-h day/night cycle at no more than five adult animals per cage. Animals were maintained on the C57BL/6J background, and newly received or generated transgenic lines were backcrossed to C57BL/6J. Experimental animals were heterozygous for the recombinase transgenes and the reporter transgenes. Transgenic lines used in this study are summarized in **Extended Data Table 5**. Standard tamoxifen treatment for CreER lines included a single dose of tamoxifen (40 μl of 50 mg ml–1) dissolved in corn oil and administered via oral gavage at postnatal day (P)10–14. Tamoxifen treatment for *Nkx2.1-CreERT2;Ai14* was performed at embryonic day (E)17 (oral gavage of the dam at 1 mg per 10 g of body weight), pups were delivered by cesarean section at E19 and then fostered. *Cux2-CreERT2;Ai14* mice received tamoxifen treatment at P35 ± 5 for five consecutive days. Trimethoprim was administered to animals containing *Ctgf-2A-dgCre* by oral gavage at P40 ± 5 for three consecutive days (0.015 ml per g of body weight using 20 mg ml–1 trimethoprim solution). *Ndnf-IRES2-dgCre* animals did not receive trimethoprim induction, since the baseline dgCre activity (without trimethoprim) was sufficient to label the cells with the *Ai14* reporter (Tasic et al., 2016). We excluded any animals with anophthalmia or microphthalmia.

### Generation of transgenic mice (*Penk-IRES2-Cre-neo, Slc17a8-IRES2-Cre, Vipr2-IRES2-Cre*)

Vectors containing gene-specific homology arms and *IRES2-Cre-bGHpoly(A)-PGK-gb2-NEO-PGKpoly(A)* components were generated using gene synthesis (GenScript) and standard molecular cloning techniques. Targeting of the transgene cassette into the endogenous gene locus immediately downstream of the stop codon was accomplished by CRISPR/Cas9-mediated genome editing using circularized targeting vector in combination with a gene-specific guide vector (Addgene plasmid #42230). The 129S6/B6 F1 ES cell line, G4 (George et al., 2007), was used to generate all modified ES cells. Correctly targeted clones were identified using standard screening approaches (PCR, qPCR and Southern blots) and injected into blastocysts to obtain chimeras and subsequent germline transmission. Resulting mice were crossed to the *Rosa26-PhiC31o* mice (JAX Stock # 007743)(Raymond and Soriano, 2007) to delete the *PGK-NEO* selection cassette, and then backcrossed to C57BL/6J mice and maintained in the C57BL/6J background. The *PGK-NEO* cassette could not be removed from *Penk-IRES2-Cre-neo* by PhiC31o-mediated recombination.

### Stereotaxic injections

*Retrograde Labeling.* We injected AAV2-retro-EF1a-Cre (Tervo et al., 2016), SADB-rabies-dGdL-Cre, or CAV-Cre (gift of Miguel Chillon Rodrigues, Universitat Autònoma de Barcelona) (Hnasko et al., 2006) into brains of heterozygous or homozygous *Ai14* mice as previously described (Tasic et al., 2016). For ALM experiments, we also injected AAV2-retro-CAG-GFP or AAV2-retro-CAG-tdTomato (Tervo et al., 2016) into wild-type mice. Stereotaxic coordinates were obtained from Paxinos adult mouse brain atlas (Franklin, 2008)(**Extended Data Table 4**). For two VISp experiments, we injected into SCs by inserting the needle through the cerebellum at a 45°-angle in the posterior to anterior direction. TdT+ or GFP+ single cells were isolated from VISp or ALM, depending on the injection area.

### Anterograde labeling

For anterograde projection mapping, we injected AAV2/1-pCAG-FLEX-EGFP-WPRE-pA (Oh et al., 2014) into VISp or ALM of 8-12 weeks old mice. The stereotaxic injection procedure was the same as described for retrograde labeling above. In *Ctgf-T2A-dgCre* mice, TMP induction was conducted 1 week after AAV injection for 3 consecutive days. Mice were sacrificed and brains perfused after 21 days (or 28 days in the case of *Ctgf-T2A-dgCre)* after AAV injection, and brains were imaged using TissueCyte 1000 system as described previously (Oh et al., 2014). Experiments can be viewed interactively on the Allen Institute data portal at http://connectivity.brain-map.org/.

### Single-cell isolation

We isolated single cells as previously described (Hempel et al., 2007; Sugino et al., 2006; Tasic et al., 2016) with modifications below. We mostly employed layer-enriching dissections, with focus on a single layer. Broader dissections (no layer enrichment or multiple layers combined) were employed for lines which label small numbers of cells, in order to facilitate isolation of sufficient number of cells. We updated our ACSF formulation to consist of CaCl_2_ (0.5 mM), glucose (25 mM), HCl (96 mM), HEPES (20 mM), MgSO_4_ (10 mM), NaH_2_PO_4_ (1.25 mM), myo-inositol (3 mM), N-acetylcysteine (12 mM), NMDG (96 mM), KCl (2.5 mM), NaHCO_3_ (25 mM), sodium L-ascorbate (5 mM), sodium pyruvate (3 mM), taurine (0.01 mM), thiourea (2 mM), and bubbled with carbogen gas (95% O_2_ and 5% CO_2_). For samples collected after 12/16/2016, the ACSF formulation also included trehalose (13.2 mM). Mice were anesthetized with isoflurane and perfused with cold carbogen-bubbled ACSF. The brain was dissected, submerged in ACSF, embedded in 2% agarose, and sliced into 250-μm coronal sections on a compresstome (Precisionary). Enzymatic digestion, trituration into single cell suspension, and fluorescence-activated cell sorting (FACS) of single cells were carried out as previously described (Tasic et al., 2016). Cells were sorted into 8-well strips containing lysis buffer from SMART-Seq v4 kit (see below) with RNase inhibitor (0.17 U/μ1), immediately frozen on dry ice, and stored at -80°C.

### cDNA amplification and library construction

We used the SMART-Seq v4 Ultra Low Input RNA Kit for Sequencing (Takara Cat#634894) to reverse transcribe poly(A) RNA and amplify cDNA according to the manufacturer’s instructions. We performed reverse transcription and cDNA amplification for 18 PCR cycles in 8-well strips, in sets of 12-24 strips at a time. At least 1 control strip was used per amplification set, which contained 4 wells without cells and 4 wells with 10 pg control RNA. Control RNA was either Mouse Whole Brain Total RNA (Zyagen, MR-201) or control RNA provided in the SMART-Seq v4 kit. All samples proceeded through NexteraXT DNA Library Preparation (Illumina FC-131-1096) using NexteraXT Index Kit V2 Set A (FC-131-2001). NexteraXT DNA Library prep was performed according to manufactorer’s instructions except that the volumes of all reagents including cDNA input were decreased to 0.4× or 0.5× by volume. Details are available in Documentation on the Allen Institute data portal at: http://celltypes.brain-map.org/.

### Sequencing data processing and QC

50-base pair paired-end reads were aligned to GRCm38 (mm10) using a RefSeq annotation gff file retrieved from NCBI on 01/18/2016 (https://www.ncbi.nlm.nih.gov/genome/annotation_euk/all/). Sequence alignment was performed using STAR v2.5.3 (Dobin et al., 2013) in twopassMode. PCR duplicates were masked and removed using STAR option “bamRemoveDuplicates”. Only uniquely aligned reads were used for gene quantification. Gene counts were computed using the R GenomicAlignments package (Lawrence et al., 2013) sumarizeOverlaps function using “IntersectionNotEmpty”. Cells with fewer than 2,000 detected genes (count > 0) and less than 30% of reads mapped to the transcriptome were removed. Doublets were removed by first classifying cells into broad classes of excitatory, inhibitory, and non-neuronal based on known markers. For each class, we selected highly specific genes that are only present in this class against all other classes, and computed the eigengene (the first principle component based on the given gene set). Eigengenes for all classes were normalized in the range of 0 and 1, and we removed any cells that scored > 0.2 in more than one class.

### Mapping reads to synthetic constructs

We mapped all non-genome-mapped reads to sequences in **Extended Data Table 6**. To avoid ambiguous counting due to stretches of sequence identity, we designated unique regions within these sequences to count mRNAs of interest. We counted only reads for which at least one of the paired ends had an overlap with the unique regions of at least 10 bp.

### Clustering

Cells that passed QC criteria were clustered using an in-house developed iterative clustering R package **iclust**, described partially in previous studies (Tasic et al., 2016; Yao et al., 2017), with new adaptions for large number of cells and improved robustness (this package will be available on Github). In brief, cells were grouped into very broad categories using known markers, then clustered using high variance gene selection, dimensionality reduction, dimension filtering, and Jaccard-Louvain or hierarchical (Ward) clustering. This process was repeated within each resulting cluster until no more child clusters met differential gene expression or cluster size termination criteria. The entire clustering procedure was repeated 100 times using 80% of all cells sampled at random, and the frequency with which cells co-cluster was used to generate a final set of clusters, again subject to differential gene expression and cluster size termination criteria. A workflow diagram for this approach is presented in **Extended Data Fig. 2**. Below, we provide more details for the analysis carried out at each iteration of clustering:

1. **Selection of high variance genes.** We first removed predicted gene models (gene names that start with Gm), genes from the mitochondrial chromosome, and ribosomal genes, as well as genes that are detected in fewer than 4 cells. To choose high variance genes, we used gene counts from each cell to fit a Loess regression curve between average scaled gene counts and dispersion (variance divided by mean). The regression residuals were then fit to a normal distribution based on 25% and 75% quantiles to calculate *p*-values and adjusted *p*-values (using Holm’s method), representing the probability that each gene had higher than expected variance. Genes were ranked by *p*-value, and the top 4,000 genes were used for dimensionality reduction.
2. **Dimensionality reduction.** The 4,000 most variable genes selected in Step 1 above were used as input for WGCNA to identify gene modules. To determine the discriminative power of each module, we used the genes in each module to cluster the cells into two clusters using Jaccard-Louvain clustering (for > 4,000 cells, (Shekhar et al., 2016)) or a combination of K-means and Ward’s hierarchical clustering (for < 4,000 cells). After dividing the cells into two clusters, we computed differential gene expression between the two clusters using the limma package for R (Ritchie et al., 2015). Differential expression (DE) gene analysis used all expressed genes, not just module-associated genes. We then computed the deScore (differential expression score), defined as the sum of −log10(adjusted P-value) of all DE genes. For deScore calculations, the maximum value each gene was allowed to contribute was 20. Only modules with deScore greater than 150 were selected for use in downstream analysis, and module eigengenes were computed for selected modules as reduced dimensions.
3. **Dimension filtering.** In previous studies (Tasic et al., 2016; Yao et al., 2017), we curated a list of gene modules that corresponded to technical biases, such the genes correlated with number of detected genes or % of reads mapping to mRNA, genes correlated with sequencing batch, and sex specific genes, which we would like to mask in downstream analysis. We computed module eigengenes for these technical bias modules, then computed the correlation between the reduced dimensions from Step 2. We removed dimensions with maximum Pearson correlation with any technical bias module greater than 0.7.
4. **Initial clustering.** For clustering, we applied either the Jaccard-Louvain method (Shekhar et al., 2016) (for > 4,000 cells), or Ward’s method (for <= 4,000 cells). The Jaccard-Louvain algorithm was applied by computing k-nearest-neighbors (k = 15) for each cell, then constructing a cell-cell similarity matrix using a Jaccard index based on the number of shared neighbors between every pair of cells. Jaccard similarity index was then used for clustering using the Louvain algorithm. This algorithm scales very well with large data sets, but has been proven to have resolution limit (Fortunato and Barthelemy, 2007), and small clusters tend to be missed. Therefore, as a complementary approach, we applied Ward’s minimum variance method for hierarchical clustering when fewer than 4000 cells were to be clustered. The initial number of clusters was set at twice the number of reduced dimensions from step 3.
5. **Cluster merging.** To make sure the resulting clusters all have distinguishable transcriptomic signatures, we defined DE genes between every cluster and their top two nearest neighbors in the reduced dimension space (using Euclidean distance if there were 1 or 2 dimensions, or 1 minus Pearson correlation for more dimensions). A pair of clusters was considered separable if the deScore (described in Step 2) for all DE genes was greater than 150. If a cluster did not pass this criterion, it was merged with the nearest cluster, then nearest neighbor and DE gene scores were recomputed using the merged clusters. Clusters with fewer than 4 cells were also merged with their nearest neighbors. This iterative merging process was repeated until all remaining clusters were separable, and have minimal cluster size. **Steps 1-5** were repeated for each resulting cluster until no further partitions were found.
6. **Defining consensus clusters.** To determine the robustness of the clustering results, the entire clustering procedure was repeated 100 times using 80% of all cells sampled at random to generate the frequency matrix for every pair of cells to co-cluster. We inferred the consensus clusters by iteratively splitting the co-clustering matrix. In any given step, we used co-clustering matrix as the similarity matrix and clustered using either Louvain (>= 4000 cells) or Ward’s algorithm (< 4000 cells). We defined *N*_k,l_ as the average probabilities of cells within cluster *k* to co-cluster with cells within cluster *l.* We merged clusters *k, l* if *N*_k,l_ > max(*N*_k,k_,*N*_1,1_) – 0.25. We merged remaining clusters based on DE genes as described in step 5 using deScore threshold of 150. We noticed that for some closely related clusters, DE genes were expressed only in one cluster, but not the other. We merged pairs of clusters that had a deScore less than 50 in either direction.
7. **Cluster Refinement.** For each cell *i*, we computed the average probability that it co-clusters with cells in each cluster *k* as *M_i,k_*, and we reassigned every cell *i* to the cluster *k* with maximum *M_i,k_* We repeated this process until convergence.
8. **Exclusion of outlier clusters.** After defining consensus clusters, we examined our clustering results to identify outlier clusters that are likely to be due technical problems.

### Assigning core and intermediate cells

In our previous study, we applied a random forest classifier to confirm our cluster assignments, and to define core and intermediate cells (Tasic et al., 2016). We found that random forest classification penalized small clusters, so we used a nearest-centroid classifier, which assigns a cell to the cluster whose centroid is the closest (with the highest correlation) to the cell. Here, the cluster centroid is defined as the median expression of 4015 differential cell type markers. To define core vs. intermediate cells, we performed 5-fold cross-validation 100 times: in each round, the cells were randomly partitioned into 5 groups, and cells in each group of 20% of the cells were classified by a nearest-centroid classifier trained using the other 80% of the cells. A cell classified to the same cluster more than 90 times was defined as a core cell, the others were designated intermediate cells. We define 19,195 core cells and 2,554 intermediate cells, which in most cases classify to only 2 clusters (2,542 out of 2,554; 99.5%).

### Correspondence between VISp and ALM excitatory clusters

To establish correspondence in both directions, we classified VISp excitatory cells using ALM clusters as training data, as well as, classified ALM excitatory cells using VISp clusters as training data. In both cases, we trained the nearest centroid classifier based on the common set of excitatory markers (pool of top 50 DE genes in each direction between excitatory clusters within VISp and ALM, respectively) shared by both regions, and calculated the fraction of cells in each VISp clusters that mapped to each of the ALM clusters, and vice versa. For each cell, we computed the correlation score of the best mapping cluster, and transformed the correlation scores into Z scores. If the average Z score of cells from one cluster mapped to another cluster in the other region was below −1.64 (roughly 5% confidence interval), this cluster was considered to be unique to one region, with no corresponding cluster in the other region.

### Assessing correspondence to Paul 2017 dataset

We mapped cells from Paul 2017 dataset to our GABAergic clusters using the nearest centroid classifier based on inhibitory markers from the current dataset that were also detected in Paul 2017 dataset (expression >0). To take into account major differences in profiling platforms we repeat classification 100 times, each time using 80% of randomly sampled markers, and computed the probabilities for every cell to map to every reference cluster.

### Assessing correspondence to Cadwell 2016 Patch-seq dataset

We mapped cells from the ArrayExpress accession E-MTAB-4092 dataset (Cadwell et al., 2016) to our VISp clusters using the nearest centroid classifier based on 3,372 of 3,406 VISp markers that were also detected in the Cadwell dataset (expression >0). To account for differences in Patch-seq and benchmark dataset (based on FACS-isolated cells) data quality, we repeated classifications 100 times, each time using 80% of randomly sampled markers, and computed the fraction of times every cell mapped to every reference cluster. Cells mapped to clusters with probabilities < 80% were mapped to the parent nodes of the mapped clusters within the cell type hierarchy, until aggregated confidence at the parent node was > 80%.

### Defining differentially expressed genes

Differentially expressed (DE) genes were detected using the R package limma v3.30.13 (Ritchie et al., 2015) using log2(CPM+1) of expression values. DE genes were defined as genes with > 2-fold change and adjusted pvalue < 0.01. We also required DE genes to have a relatively bimodal expression pattern, expressed predominately in one cluster relative to the other. To do that, we computed *P_i,j_* as the fraction of cells in cluster *j* expressing gene *i* with CPM ≥ 1, and required up-regulated genes *i* in cluster *c*_1_ relative to *c*_2_ to have *P_i,c_*_1_ > 0.4, and (*P_i,c_*_1_ — *P_i,c_*_2_)/max((*P_i,c_*_1_,*P_i,c_*_2_) > 0.7. Due to significant computation overhead, we only used the “voom” mode, which increases the accuracy of limma on RNA-seq datasets, to calculate DE genes for the final clusters, and utilized limma’s default mode for cluster merging steps (Steps 5 and 6, above).

### Measures of heterogeneity within VISp-L4-Rspo1 and between VISp-L4-Rspo1 and related clusters

To explore the heterogeneity of the VISp-L4-Rspo1 cluster, which corresponds to three separate cell types in our previous study ((Tasic et al., 2016) and **Extended Data Fig.7**), we first removed the QC-dependent gene expression signatures by regressing the expression of each gene against the QC index, defined as the ratio of the fraction of the reads mapped to non-exonic regions over reads mapped to exonic regions. This QC-dependent signature is particularly high for L4 cells, and is confounded with other transcriptomic signatures. After normalization, we performed WGCNA to find co-expressed gene modules within cells from VISp-L4-Rspo1. We found that the eigengene for the top gene module within VISp L4 Rspo1 corresponds to the gradient that drove separation of L4 subtypes previously (Tasic et al., 2016). We then took the 50 cells that were most correlated with each end of the eigengene-defined gradient, and used their gene expression data to train a random forest classifier. The classification probabilities by random forest strongly correlated with the gradient eigengene (**Extended Data Fig. 7e**). The distribution of the classification probabilities was compared with a uniform distribution using a Kolmogorov-Smirnov test (KS test), which gave a modest p-value when used to test cells from within the VISp-L4-Rspo1 cluster (6.69×10^-5^). We repeated the same test between VISp-L4-Rspo1 and the neighboring VISp-L5-Endou cluster. The eigengene for this comparison was defined as the first principle component of the top 50 DE genes in both directions. In this case, the random forest classfication probabilities deviated from uniform distribution more significantly than when cells within VISp-L4-Rspo1 alone were used (KS Test p-value = 2.9×10^-9^). When cells in the VISp-L4-Rspo1 cluster were compared with the more distant VISp-L5-Batf3 cluster, the separation is clear: classification probabilities have a bimodal distribution.

### RNA FISH by RNAScope

We performed RNA FISH using RNAScope Multiplex Fluorescent v1 and v2 kits (Advanced Cell Diagnostics, Newark, CA) according to the manufacturer’s protocol. We used fresh frozen sections, which we prepared by dissecting fresh brains, embedding the brains in OCT, and storing OCT blocks at −80 °C. 10-μm coronal sections were cut using a cryostat and collected on SuperFrost slides. Sections were allowed to dry for 30 min at −20 °C in the cryostat chamber, placed into pre-chilled plastic slide boxes, wrapped in a zipped plastic bag, and stored at -80 °C. Slides were used within one week. Nuclei were labeled by DAPI and nuclear signal was used to segment cells in images. We imaged mounted sections at 40X on a confocal microscope (Leica SP8). Maximum projections of z-stacks (1-μm intervals) were processed using CellProfiler [www.cellprofiler.org] (Lamprecht et al., 2007) to identify nuclei, quantify the number of fluorescent spots, and assign fluorescent spots to each nucleus. Diagrmas were generated by plotting each cell as a dot at its nucleus position and colored it according to the quantified fluorescent labeling.

### Immunohistochemistry

Mice were perfused with 4% paraformaldehyde (PFA). Brains were dissected and post-fixed with 4% PFA at room temperature for 3-6 h followed by overnight at 4 °C. Brains were rinsed with PBS and cryoprotected in 10% sucrose (w/v) in PBS with 0.1% sodium-azide overnight at 4 °C. 100-μm coronal slices were sectioned on a microtome (Leica #SM2010R), washed with PBS, blocked with 5% normal donkey serum in PBS and 0.3% Triton X-100 (PBST) for 1 h, and stained with rabbit anti-dsRed (1:1000, Clontech #632496) and goat anti-Pvalb (1:1000, Swant #PVG-213) overnight at room temperature. Sections were washed in three times in PBST and incubated with anti-rabbit Alexa 594 (1:500, Jackson ImmunoResearch #711-585-152) and anti-goat Alexa 488 (1:500, Jackson ImmunoResearch #705-605-147) for 4 h at room temperature. Sections were washed three times with PBST and stained with 5 μM DAPI in PBS for 20 min. After washing in PBST, sections were mounted onto slides, allowed to dry, rehydrated in PBS, dipped in water and cover slipped with Fluoromount G (SouthernBiotech #0100-01) mounting medium.

### Data analysis software and visualization tools

Analysis and visualization of transcriptomic data wereperformed using R v3.3.0 and greater, as well as the following R packages: cowplot v0.8.0, dendextend v1.5.2, dplyr v0.7.4, feather v0.3.1, GenomicAlignments v1.12.2, ggbeeswarm v0.6.0, ggplot2 v2.2.1, ggrepel v0.7.0, googlesheets v0.2.2, Hmisc v4.0-3, limma v3.30.13, Matrix v1.2-11, matrixStats v0.52.2, purr v0.2.3, pvclust v2.0-0, reshape2 v1.4.2, Rtsne v0.13, Seurat v2.1.0, viridis v0.4.0, WGCNA v1.61, and xlsx v0.5.7.

### Data, reagent, and code availability

Single cell transcriptomic data is in the process of deposition to GEO. Full metadata for all samples are available in **Extended Data Table 8**. Newly generated mouse lines have been deposited to the Jackson Laboratory: *Vipr2-IRES2-Cre* (JAX stock number pending), *Slc17a8-IRES2-Cre* (JAX stock number pending), *Penk-IRES2-Cre-neo* (JAX stock number 025112). Code used for analysis and figures will be available from GitHub at github.com/AllenInstitute/ (code deposit in progress). An R package for the iterative clustering method utilized in this analysis will be available on GitHub at github.com/AllenInstitute/ (code deposit in progress).

